# *Shigella sonnei* infection of zebrafish reveals that O-antigen mediates neutrophil tolerance and dysentery incidence

**DOI:** 10.1101/719781

**Authors:** Vincenzo Torraca, Myrsini Kaforou, Jayne Watson, Gina M. Duggan, Hazel Guerrero-Gutierrez, Sina Krokowski, Michael Hollinshead, Thomas B. Clarke, Rafal J. Mostowy, Gillian S. Tomlinson, Vanessa Sancho-Shimizu, Abigail Clements, Serge Mostowy

**Affiliations:** Section of Microbiology, MRC Centre for Molecular Bacteriology and Infection, Imperial College London, London, UK; Department of Infection Biology, London School of Hygiene & Tropical Medicine, London, UK; Department of Paediatrics, Division of Medicine, Imperial College London, London, UK; Faculty of Natural Sciences, Department of Life Sciences, MRC Centre for Molecular Bacteriology and Infection, Imperial College London, London, UK; Division of Virology, Department of Pathology, Cambridge University, Cambridge, UK; Malopolska Centre of Biotechnology, Jagiellonian University, Krakow, Poland; Faculty of Medicine, School of Public Health, Imperial College London, London, UK; Division of Infection and Immunity, University College London, London, UK; Department of Virology, Division of Medicine, Imperial College London, London, UK

**Author notes:** Corresponding author: Prof. Serge Mostowy, Department of Infection Biology, London School of Hygiene & Tropical Medicine, Keppel St, London WC1E 7HT UK. **Author contributions:** V.T. and S.M. designed the research and analysed the data. V.T., G.M.D., H.G.G., S.K. and M.H. performed experiments. J.W. and A.C. contributed bacterial strains. M.K., G.S.T. and R.J.M. provided supervision and assistance with RNAseq data analysis and presentation. T.B.C. and V.S.S. provided supervision and assistance with neutrophil infection experiments. V.T. and S.M. wrote the manuscript with input from all the authors.

**Keywords:** neutrophils, O-antigen, *Shigella flexneri*, *Shigella sonnei*, zebrafish

## Abstract

*Shigella flexneri* is historically regarded as the primary agent of bacillary dysentery, yet the closely-related *Shigella sonnei* is replacing *S. flexneri*, especially in developing countries. The underlying reasons for this dramatic shift are mostly unknown. Using a zebrafish (*Danio rerio*) model of *Shigella* infection, we discover that *S. sonnei* is more virulent than *S. flexneri in vivo*. Whole animal dual-RNAseq and testing of bacterial mutants suggest that *S. sonnei* virulence depends on its O-antigen oligosaccharide (which is unique among *Shigella* species). We show *in vivo* using zebrafish and *ex vivo* using human neutrophils that *S. sonnei* O-antigen can mediate neutrophil tolerance. Consistent with this, we demonstrate that O-antigen enables *S. sonnei* to resist phagolysosome acidification and promotes neutrophil cell death. Chemical inhibition or promotion of phagolysosome maturation respectively decreases and increases neutrophil control of *S. sonnei* and zebrafish survival. Strikingly, larvae primed with a sublethal dose of *S. sonnei* are protected against a secondary lethal dose of *S. sonnei* in an O-antigen-dependent manner, indicating that exposure to O-antigen can train the innate immune system against *S. sonnei*. Collectively, these findings reveal O-antigen as an important therapeutic target against bacillary dysentery, and may explain the rapidly increasing *S. sonnei* burden in developing countries.

**Author Summary:** *Shigella sonnei* is predominantly responsible for dysentery in developed countries, and is replacing *Shigella flexneri* in areas undergoing economic development and improvements in water quality. Using *Shigella* infection of zebrafish (*in vivo*) and human neutrophils (*in vitro*), we discover that *S. sonnei* is more virulent than *S. flexneri* because of neutrophil tolerance mediated by its O-antigen oligosaccharide acquired from the environmental bacteria *Plesiomonas shigelloides*. To inspire new approaches for *S. sonnei* control, we show that increased phagolysosomal acidification or innate immune training can promote *S. sonnei* clearance by neutrophils *in vivo*. These findings have major implications for our evolutionary understanding of *Shigella*, and may explain why exposure to *P. shigelloides* in low and middle-income countries (LMICs) can protect against dysentery incidence.

## Introduction

*Shigella* is the causative agent of bacillary dysentery (also called shigellosis), resulting from invasion of the intestinal epithelium and leading to ∼164,000 deaths annually [1, 2]. *Shigella* is also recognised by the World Health Organization as a priority pathogen exhibiting antimicrobial resistance [3, 4]. The emergence of multidrug resistant bacteria and the lack of effective vaccines has resulted in a desperate need to understand *Shigella* pathogenesis and identify new approaches for infection control. In the lab, infection with *Shigella flexneri* has been a valuable discovery tool in the field of innate immunity, helping to illuminate the role of neutrophil extracellular traps (NETs) [5], nucleotide-binding oligomerisation domain (NOD)-like receptors (NLRs) [6], bacterial autophagy [7], interferon-inducible guanylate-binding proteins (GBPs) [8, 9] and septin-mediated cell-autonomous immunity [10, 11] in host defence.

The genus *Shigella* comprises four different species (*S. flexneri*, *S. sonnei*, *S. boydii*, *S. dysenteriae*), although DNA sequencing suggests they evolved from convergent evolution of different founders [12]. The most recent strains of *S. flexneri* emerged from *Escherichia coli* >35,000 years ago [12], while *S. sonnei* (a monoclonal strain) emerged from *E. coli* in central Europe ∼500 years ago [13]. *S. flexneri* is historically regarded as the primary agent of dysentery worldwide, yet *S. sonnei* has recently become the most prevalent cause of dysentery in developing countries (i.e., areas undergoing economic development and improvements in water quality) [14, 15]. Reasons for this dramatic shift are mostly unknown. Hypotheses include improved water sanitisation leading to reduced cross-immunisation by *Plesiomonas shigelloides* (which carries an O-antigen oligosaccharide identical to *S. sonnei*) [15, 16], as well as a type VI secretion system (T6SS)-mediated competitive advantage that *S. sonnei* exerts over *S. flexneri* and the Gram-negative gut microbiome for niche occupancy [17].

Except for non-human primates, there is no mammalian model that fully recapitulates human shigellosis. The zebrafish model is increasingly being used to study human bacterial pathogens *in vivo*, including *S. flexneri* [18, 19]. The major pathogenic events that lead to shigellosis in humans (i.e., macrophage cell death, invasion and multiplication within epithelial cells, cell-to-cell spread, inflammatory destruction of the host epithelium) are recapitulated in a zebrafish model of *S. flexneri* infection [20]. Exploiting the optical accessibility of zebrafish larvae, it is possible to spatio-temporally examine the development, coordination and resolution of the innate immune response to *S. flexneri in vivo*. As a result, *S. flexneri*-zebrafish infection has been useful to illuminate key roles for bacterial autophagy [21], bacterial predation [22], inflammation [23] and trained innate immunity [24] in host defence *in vivo*.

How *S. sonnei* infection differs from *S. flexneri* infection is poorly understood, yet clinical management of both infections is the same. Here, we develop a *S. sonnei*-zebrafish infection model and discover that *S. sonnei* is more virulent than *S. flexneri in vivo* because of neutrophil tolerance mediated by its O-antigen. We show that increased phagolysosomal acidification or innate immune training can promote *S. sonnei* clearance by neutrophils *in vivo*. These results may inspire new approaches for *S. sonnei* control.

## Results

### *S. sonnei* is more virulent than *S. flexneri* in a zebrafish infection model

To compare the virulence of *S. flexneri* and *S. sonnei in vivo*, we injected *S. flexneri* M90T or *S. sonnei* 53G in the hindbrain ventricle (HBV) of zebrafish larvae at 3 days post-fertilisation (dpf). Unexpectedly, *S. sonnei* led to significantly more zebrafish death and higher bacterial burden, as compared to *S. flexneri* (**Fig. 1A,B**). The majority of larvae inoculated with <600 CFU of *S. sonnei* survive, whereas the majority of larvae inoculated with >1500 CFU of *S. sonnei* die by 72 hpi (**Fig. S1A,B**). In agreement with being more virulent, *S. sonnei* infected larvae have significantly increased expression of key inflammatory markers at 6 and 24 hpi, as compared to *S. flexneri* infected larvae (**Fig. 1C,D**). Moreover, *S. sonnei*, unlike *S. flexneri*, disseminates out of the HBV into the neuronal tube and bloodstream (**Fig. 1E,F, Fig. S1C-F)**. The increased virulence of *S. sonnei* is also observed using an intravenous route of infection (**Fig. S1G-H**), using human clinical isolates of bacteria (**Fig. S1I-J**) and when infected larvae are incubated at 28.5°C, 32.5°C or 37°C (**Fig. 1A,B, Fig. S1K-N**). Therefore, we used *S. sonnei* 53G infection of the HBV (incubated at 28.5°C, the standard temperature for zebrafish maintenance) for the rest of our study. Taken together, using multiple infection routes, bacterial strains and temperatures, these data show for the first time that *S. sonnei* is significantly more virulent than *S. flexneri in vivo*.

**Figure 1.**
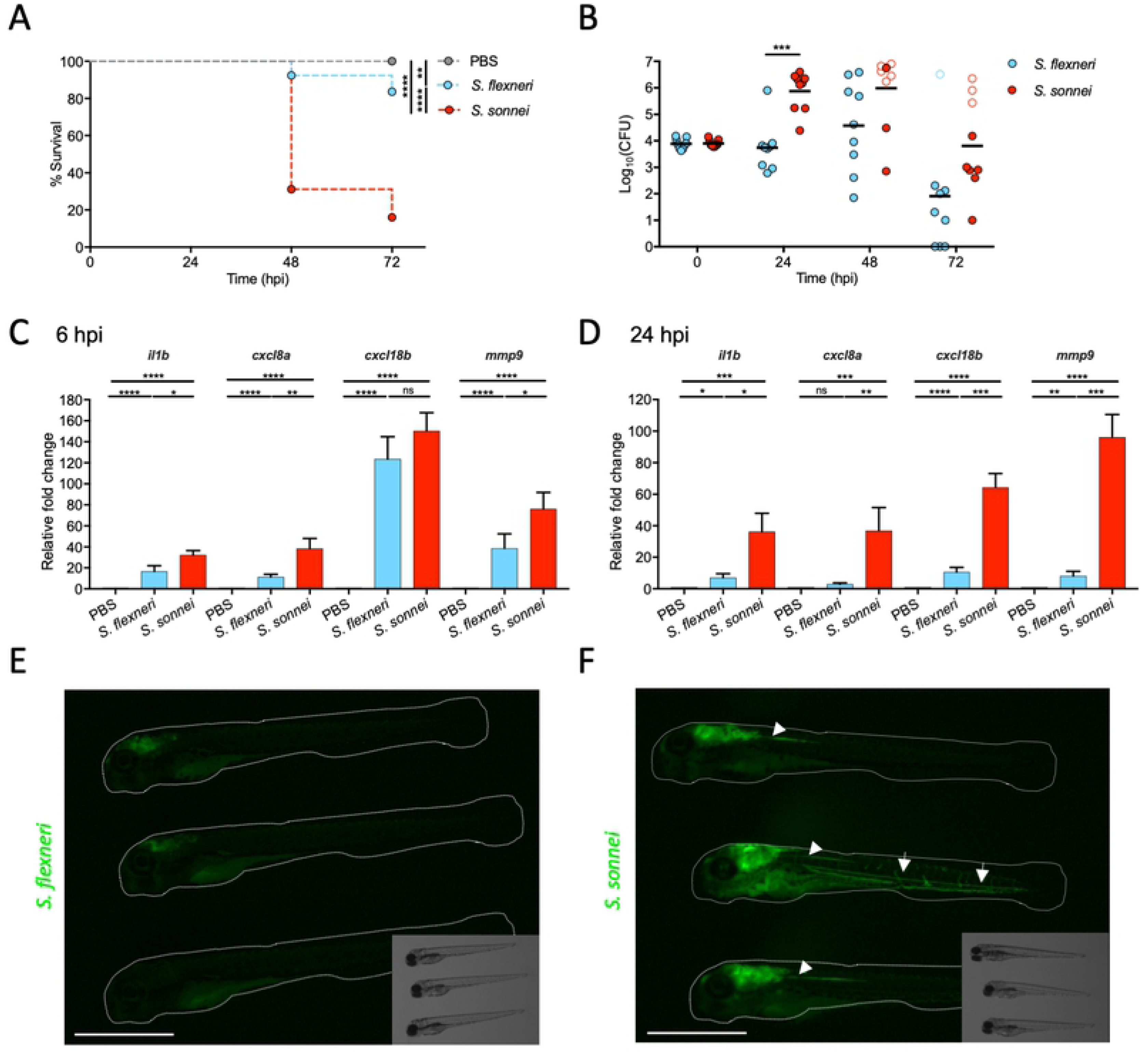
*S. sonnei* is more virulent than *S. flexneri* in a zebrafish infection model. **A,B. *S. sonnei* is more virulent than *S. flexneri in vivo***. Survival curves (A) and Log_10_-transformed CFU counts (B) of larvae injected in the hindbrain ventricle (HBV) with PBS (grey), *S. flexneri* (blue) or *S. sonnei* (red). Experiments are cumulative of 3 biological replicates. In B, full symbols represent live larvae and empty symbols represent larvae that at the plating timepoint had died within the last 16 hours. Statistics: Log-rank (Mantel-Cox) test (A); unpaired t-test on Log_10_-transformed values (B); **p<0.0021; ***p<0.0002; ****p< 0.0001. **C,D. *S. sonnei* elicits a stronger inflammatory signature than *S. flexneri in vivo***. Quantitative real time PCR for representative inflammatory markers were performed on pools of 20 HBV injected larvae collected at 6 (C) or 24 (D) hpi with PBS (grey), *S. flexneri* (blue) or *S. sonnei* (red). Experiments are cumulative of 4 biological replicates. Statistics: one-way ANOVA with Sidak’s correction on Log_2_-transformed values; ns, non-significant; *p<0.0332 **p<0.0021; ***p<0.0002; ****p<0.0001. **E,F. *S. sonnei* can disseminate from the injection site**. Representative images of three GFP-labelled *S. flexneri*-infected (E) or S. *sonnei*-infected (F) larvae at 24 hpi. In D, arrows indicate dissemination in the blood circulation; arrowheads indicate dissemination in the neuronal tube. Scale bars = 1 mm.

### Whole animal dual-RNAseq profiling of *S. sonnei* infected larvae

The transcriptional signature of *Shigella in vivo* was mostly unknown. We performed whole animal dual-RNAseq profiling of *S. sonnei* infected larvae by isolating whole RNA at 24 hpi and mapping reads to both *Shigella* and zebrafish genomes (**Fig. 2A**). RNA isolated from Log phase (OD_600_ ∼0.6) bacterial culture grown at 28.5°C was used as baseline for the identification of differential expression in the bacterial transcriptome. RNA isolated from PBS-injected larvae was used as baseline for the identification of differential expression in the host transcriptome. Gene count data was obtained for both the host and pathogen, and statistical analysis was performed using DESeq2 (see dedicated section in Materials and Methods). Genes were considered significantly differentially expressed if Log_2_(FC) > 1 or < -1 and the adjusted p value < 0.05. Principal component analysis (PCA) was employed for *S. sonnei* and larval count data separately, and these plots confirm the clustering between biological replicates according to their state (**Fig. S2A,B**). After performing differential expression analysis between infected and control states, we found 1538 differentially expressed *S. sonnei* genes (representing ∼1/3 of the *S. sonnei* 53G genome, **Fig. 2B**, see also **Table S1** and **Fig. S2** for in-depth exploration) and 337 differentially expressed zebrafish genes (**Fig. 2C**, see also **Table S2** and **Fig. S2** for in-depth exploration). In the case of *S. sonnei*, 878 genes are significantly upregulated, including genes involved in resistance to stress (i.e. adaptation to acidic environment, metabolism, DNA damage repair; **Fig. 2D**, see also **Fig. S2E,G,J** and **Table S1**). In the case of *S. sonnei*-infected larvae, 283 host genes are significantly upregulated, including inflammatory markers previously tested by qRT-PCR (**Fig. 1C,D**) and other genes involved in innate immune signalling, granulopoiesis/neutrophil chemotaxis and inflammation (**Fig. 2E**, see also **Fig. S2F,H,I,K** and **Table S2).** Consistent with this, enrichment analysis for DNA regulatory elements identified the statistical overrepresentation of immune-related transcription factor binding sites (i.e. Rel/Rela, Nfkb2, Stats, Cepbg, Jun, Spi1; **Table S2**). Together, whole animal dual-RNAseq profiling identified novel markers of *S. sonnei* infection and zebrafish host defence, and we generated an open-access resource for their in-depth exploration (raw sequencing data will be uploaded into GEO, the NCBI repository).

**Figure 2.**
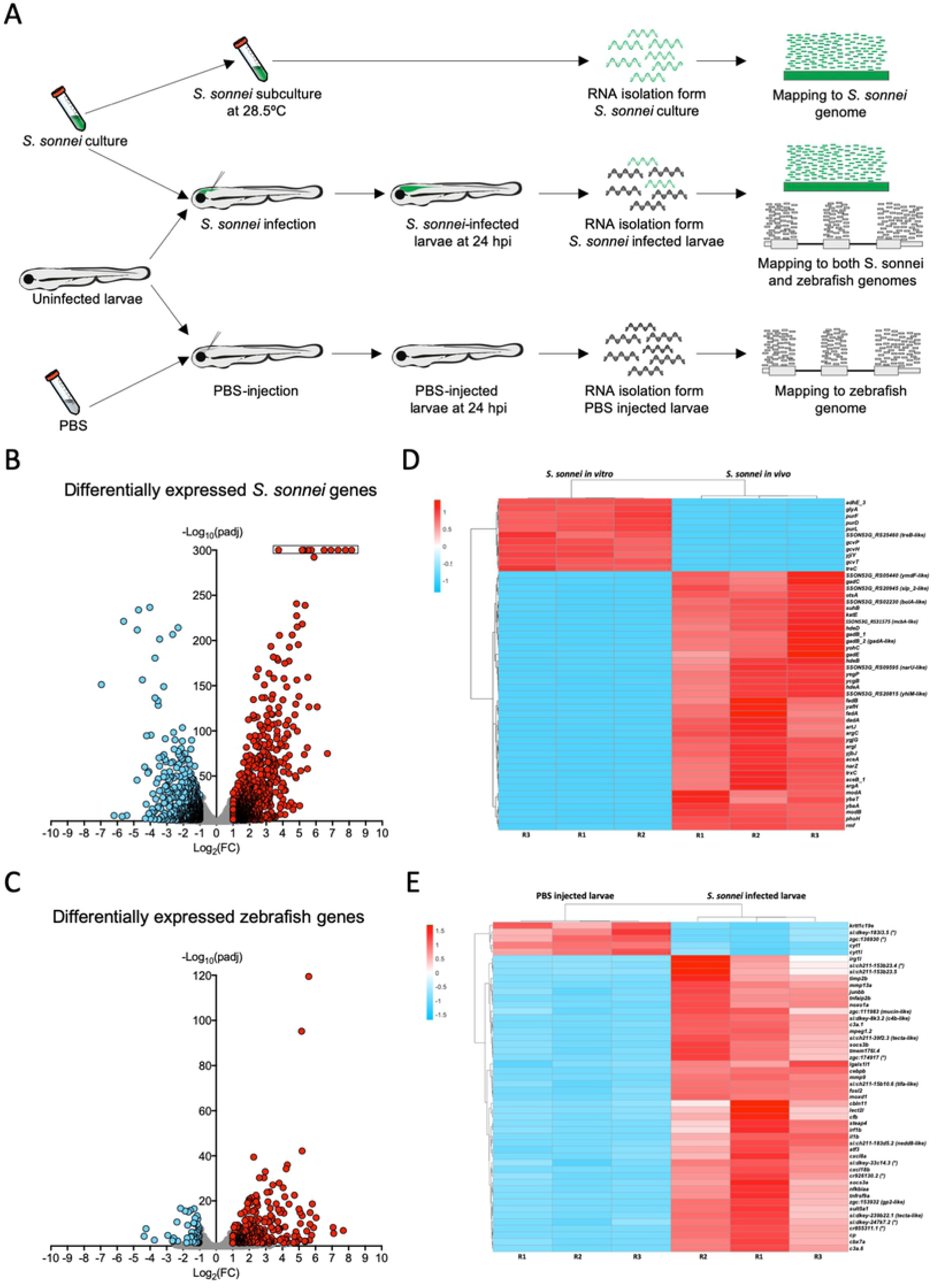
Whole animal dual-RNAseq profiling of *S. sonnei* infected larvae. **A. Workflow for dual-RNAseq processing.** 3 dpf larvae were infected with ∼7000 CFU of *S. sonnei*. Pools of infected larvae were collected for RNA isolation at 24 hpi. As a control for the bacterial transcriptome, the same cultures of *S. sonnei* were diluted 50x and subcultured at 28.5°C (same temperature at which infected larvae are maintained) until Log phase (OD_600_ ∼0.6) was reached. As a control for the zebrafish transcriptome, pools of PBS-injected larvae at 24 hpi were used. Reads from infected larvae were mapped separately to both the *S. sonnei* and zebrafish genomes, while reads from *S. sonnei in vitro* cultures and PBS injected larvae were mapped to the pathogen or host genome, respectively. **B,C. Volcano plots for bacterial and zebrafish genes during *S. sonnei* infection.** Each datapoint refers to a single gene. Non significantly differentially expressed genes are shown in grey, while significantly downregulated genes are shown in blue and significantly upregulated genes are shown in red. Plot in B refers to *S. sonnei* genes and plot in C refers to zebrafish genes. See also **Fig. S2** for additional details. Log_2_(FC) and -Log_10_(padj) coordinates were derived from data analysis with the DESeq2 package in R. In B, points enclosed in the black rectangle were computed to have a DESeq2 padj = 0. **D,E. Heatmap of the top 50 differentilly expressed bacterial and zebrafish genes during *S. sonnei* infection.** Columns represent individual biological replicates (R1, R2, R3). Heatmaps were created from counts per million (CPM) reads values, using “pheatmap” package in R. Shades of blue indicate downregulation and shades of red indicate upregulation compared to baseline. Plot in D refers to *S. sonnei* genes and plot in E refers to zebrafish genes. Gene names in brackets are inferred by manual annotations based on protein aligments performed on the Uniprot database (https://www.uniprot.org/). For genes not predicted to encode a protein, manual annotations were inferred from the Ensembl database (https://www.ensembl.org/). (*) *si:dkey-183i3.5:* thread biopolymer filament subunit alpha-like*; zgc:136930:* thread biopolymer filament subunit gamma-like; *si:ch211-153b23.4: YrdC domain-containing protein-like; zgc:174917:* phytanoyl-CoA dioxygenase domain-containing protein 1-like; *si:dkey-247k7.2:* carboxypepD reg-like domain-containing protein-like; *si:dkey-33c14.3* and *cr926130.2*: uncharacterised lincRNAs; *cr855311.1:* uncharacterised non-coding RNA. See also **Tables S1,2** for the extended gene lists and **Fig. S2** for in-depth exploration of the data.

### *S. sonnei* virulence depends on its O-antigen

*S. sonnei* encodes a T6SS and capsule which have been linked to virulence in the murine and rabbit intestine model, respectively (**Fig. S3A,B**) [17, 25]. However, when larvae are infected with a T6SS (*Δtssb*) or capsule (*Δg4c*) deficient strain, virulence is not significantly reduced as compared to wildtype (WT) bacteria (**Fig. S3C,D**). We next infected larvae with a phase II *S. sonnei* strain which lacks the pSS virulence plasmid (-*pSS S. sonnei*). Here, ∼100% of larvae survive infection (**Fig. 3A,B**). Consistent with a role in virulence, the *S. sonnei* pSS plasmid (which is unstable and frequently lost in culture [26]) is retained during zebrafish infection at 28.5°C (**Fig. S3E**). The *S. sonnei* pSS encodes a type III secretion system (T3SS) and the biosynthesis machinery for an O-antigen oligosaccharide (O-Ag) non-homologous to those encoded by other *Shigella* species (**Fig. S3F**) [27]. Surprisingly, infection with O-Ag deficient (*ΔO-Ag*) *S. sonnei*, and not T3SS deficient (*Δmxid*) *S. sonnei*, recapitulates results obtained with *S. sonnei* -*pSS* (**Fig. 3A-D**). Consistent with a role for O-Ag in *S. sonnei* virulence, dual-RNAseq profiling identified *wzzB*/SSON53G_RS12230 (a protein involved in the extension of O-Ag oligosaccharide chains) as significantly upregulated (Log_2_(FC) = 1.31, padj = 3.31·10^−25^) during zebrafish infection (**Fig. S3G**).

**Figure 3.**
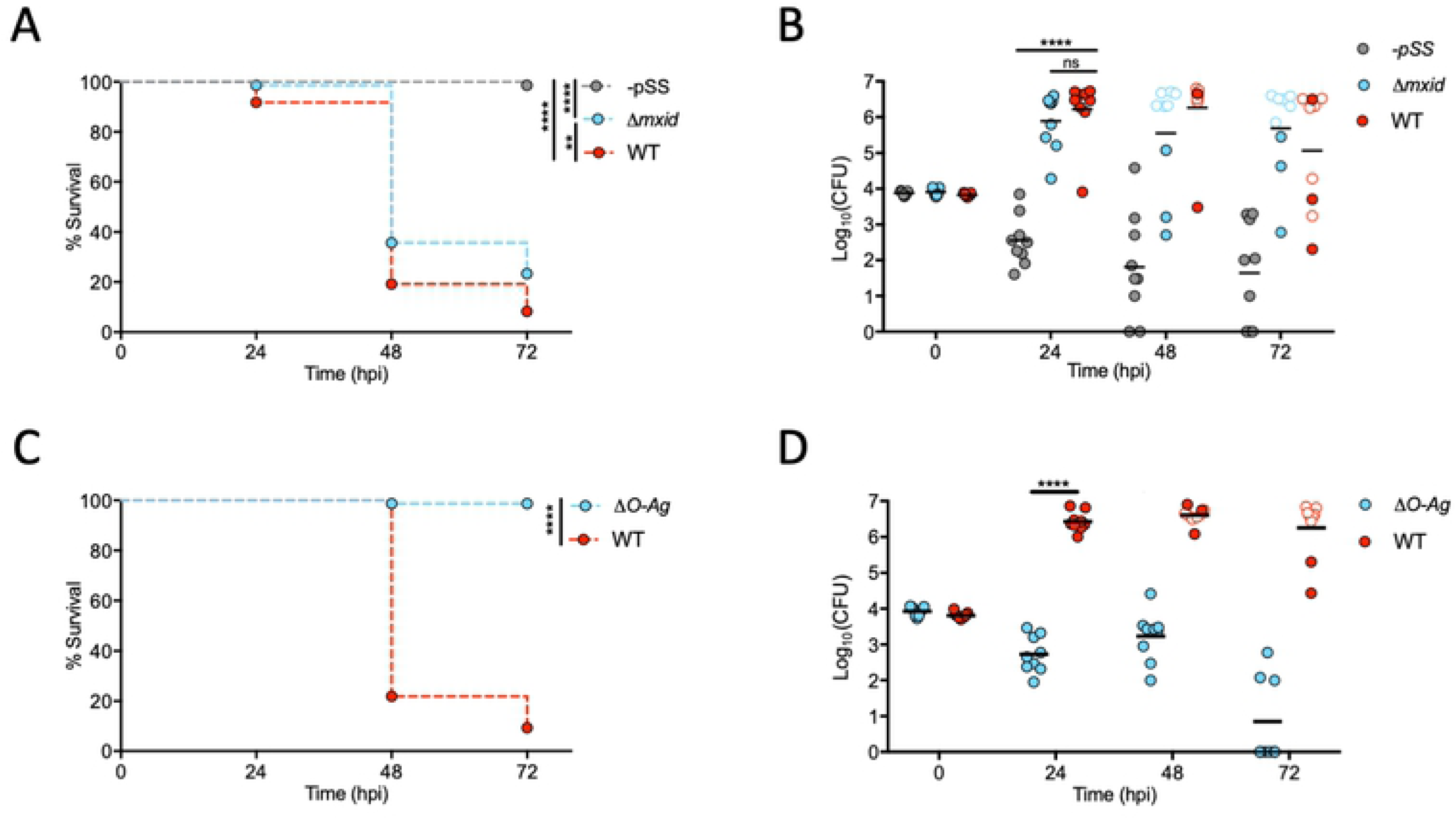
*S. sonnei* virulence depends on its O-antigen. **A,B. Virulence of *S. sonnei* depends on its virulence plasmid**. Survival curves (A) and Log_10_-transformed CFU counts (B) of larvae injected in the HBV with *S. sonnei -pSS* (grey), *Δmxid* (blue) or WT (red) strains. Experiments are cumulative of 3 biological replicates. In B, full symbols represent live larvae and empty symbols represent larvae that at the plating timepoint had died within the last 16 hours. Statistics: Log-rank (Mantel-Cox) test (A); one-way ANOVA with Sidak’s correction on Log_10_-transformed data (B); ns, non-significant; **p<0.0021; ****p<0.0001. **C,D. Virulence of *S. sonnei* depends on its O-antigen.** Survival curves (C) and Log_10_-transformed CFU counts (D) of larvae injected in the HBV with *S. sonnei ΔO-Ag* (blue) or WT (red) strains. Experiments are cumulative of 3 biological replicates. In D, full symbols represent live larvae and empty symbols represent larvae that at the plating timepoint had died within the last 16 hours. Statistics: Log-rank (Mantel-Cox) test (C); unpaired t-test on Log_10_-transformed data (B); ****p<0.0001.

### *S. sonnei* O-antigen can counteract clearance by zebrafish neutrophils

Considering that injection of *S. sonnei* induces recruitment of both macrophages and neutrophils to the HBV at 6 hpi (**Fig. 4A,B**), we tested the role of these immune cells in *S. sonnei* virulence. We used the zebrafish line *Tg(mpeg1:Gal4-FF)^gl25^/Tg(UAS-E1b:nfsB.mCherry)^c264^* in which treatment with the pro-drug Metronidazole (Mtz) results in macrophage ablation (**Fig. S4A-C**). Here, the presence or absence of macrophages does not significantly affect zebrafish survival or bacterial burden (**Fig. 4C,D, Fig. S4D,E**). In contrast, when both macrophages and neutrophils are depleted using *pu.1* morpholino oligonucleotide, we observe a significant increase in zebrafish susceptibility to *S. sonnei* and bacterial burden (**Fig. 4E,F**).

**Figure 4.**
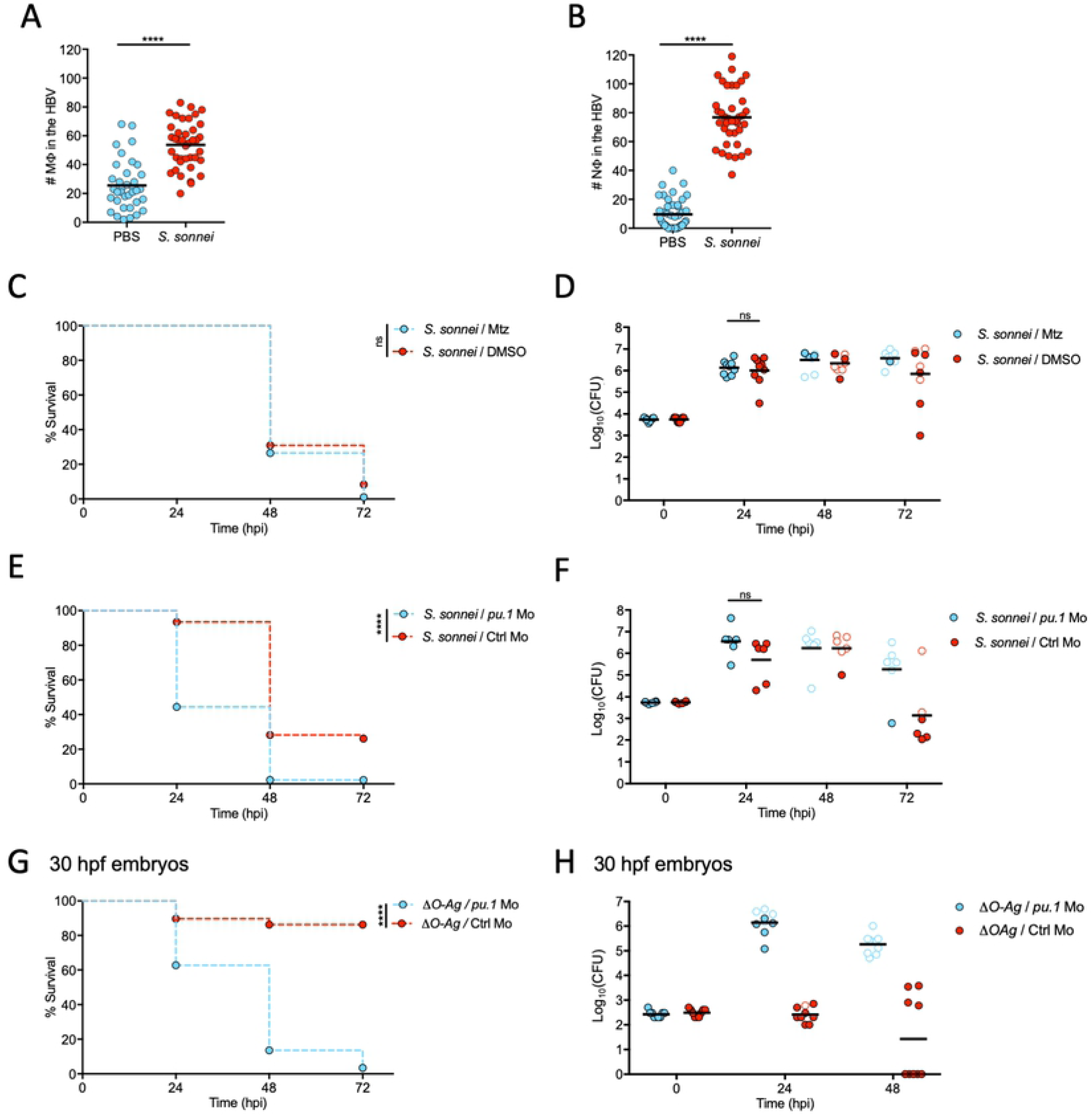
*S. sonnei* O-antigen can counteract clearance by zebrafish neutrophils. **A,B. Macrophages and neutrophils are recruited to *S. sonnei in vivo***. Larvae of the *Tg(mpeg1:Gal4-FF)^gl25^/Tg(UAS-E1b:nfsB.mCherry)^c264^* strain (labelling macrophages, A) or of the *Tg(lyz:dsRed)^nz50^* strain (labelling neutrophils, B) were injected with PBS (blue) or *S. sonne*i (red) in the HBV. Recruitment was quantified from images at 6 hpi. Experiments are cumulative of 3 biological replicates. Statistics: two-tailed Mann-Whitney test; ns, non-significant; ****p<0.0001. **C,D. Macrophage ablation does not increase susceptibility to *S. sonnei*.** Survival curves (C) and Log_10_-transformed CFU counts (D) of *Tg(mpeg1:Gal4-FF)^gl25^/Tg(UAS-E1b:nfsB.mCherry)^c264^* larvae which were treated with either Metronidazole (Mtz, macrophage ablated group, blue) or control DMSO vehicle (DMSO, red) prior to infection in the HBV with *S. sonnei*. Experiments are cumulative of 3 biological replicates. In D, full symbols represent live larvae and empty symbols represent larvae that at the plating timepoint had died within the last 16 hours. Statistics: Log-rank (Mantel-Cox) test (C); unpaired t-test on Log_10_-transformed data (D); ns, non-significant. **E,F. *pu.1* morpholino knockdown increases susceptibility to *S. sonnei*.** Survival curves (E) and Log_10_-transformed CFU counts (F) of *pu.1* morphant (blue) or control (red) larvae infected in the HBV with *S. sonnei*. Experiments are cumulative of 3 biological replicates. In F, full symbols represent live larvae and empty symbols represent larvae that at the plating timepoint had died within the last 16 hours. Statistics: Log-rank (Mantel-Cox) test (E); unpaired t-test on Log_10_-transformed data (F); ns, non-significant; ****p<0.0001. **G,H. Virulence of *ΔO-Ag S. sonnei* can be observed in *pu.1* morphants.** Survival curves (G) and Log_10_-transformed CFU counts (H) of *pu.1* morphant (blue) or control (red) larvae infected in the HBV with *ΔO-Ag S. sonnei*. To allow full ablation of immune cells by morpholino knockdown, infections were performed at 30 hours post-fertilisation (hpf). Experiments are cumulative of 3 biological replicates. In H, full symbols represent live larvae and empty symbols represent larvae that at the plating timepoint had died within the last 16 hours. Statistics: Log-rank (Mantel-Cox) test (G); unpaired t-test on Log_10_-transformed data (H); ****p<0.0001.

Considering an important role for neutrophils in *Shigella* control [21, 23], and the significant upregulation of genes involved in granulopoiesis/neutrophil chemotaxis identified by dual-RNAseq profiling (e.g., *cebpb*, *atf3*, *cxcl8a*, *cxcl18b* (**Fig. 1C,D, Fig. 2E**)), we reasoned that *ΔO-Ag S. sonnei* may be attenuated *in vivo* because of its inability to counteract neutrophil clearance. Consistent with this hypothesis, infection of *pu.1* morphants at 30 hpf (when depletion of immune cells is complete) with *ΔO-Ag S. sonnei* led to significantly increased zebrafish death and bacterial burden, compared to control morphants at the same developmental stage (**Fig. 4G,H, Fig. S4F-G**).

### *S. sonnei* can resist phagolysosome acidification and promote neutrophil cell death in an O-antigen-dependent manner

Using high resolution confocal microscopy, we observed that *S. sonnei* mostly reside within neutrophil phagosomes at 3 hpi (**Fig. 5A**). Additionally, staining of live bacteria with a pH sensitive dye (pHrodo) showed that intracellular *S. sonnei* mostly reside within acidic compartments (**Fig. 5B**). We therefore hypothesised that O-Ag may promote bacterial survival during phagolysosome acidification. To test this, we measured the growth of WT or *ΔO-Ag S. sonnei* grown in liquid culture at different pH. While the growth of both strains is similar at neutral pH = 7, WT *S. sonnei* grew significantly faster than *ΔO-Ag S. sonnei* at pH = 5 (**Fig. 5C,D, Fig. S5A,B**). Consistent with a role for O-Ag in tolerance to phagolysosome acidification, transmission electron microscopy (TEM) of zebrafish larvae at 3 hpi showed intact and dividing WT *S. sonnei* cells, versus disrupted and non-dividing *ΔO-Ag S. sonnei* cells, in neutrophil phagosomes (**Fig. 5E,F, Fig. S5C**). Moreover, only in the case of WT *S. sonnei* could we observe compromised nuclei and extranuclear chromatin in zebrafish cells harbouring infection, indicative of necrotic cell death (**Fig. S5D**).

**Figure 5.**
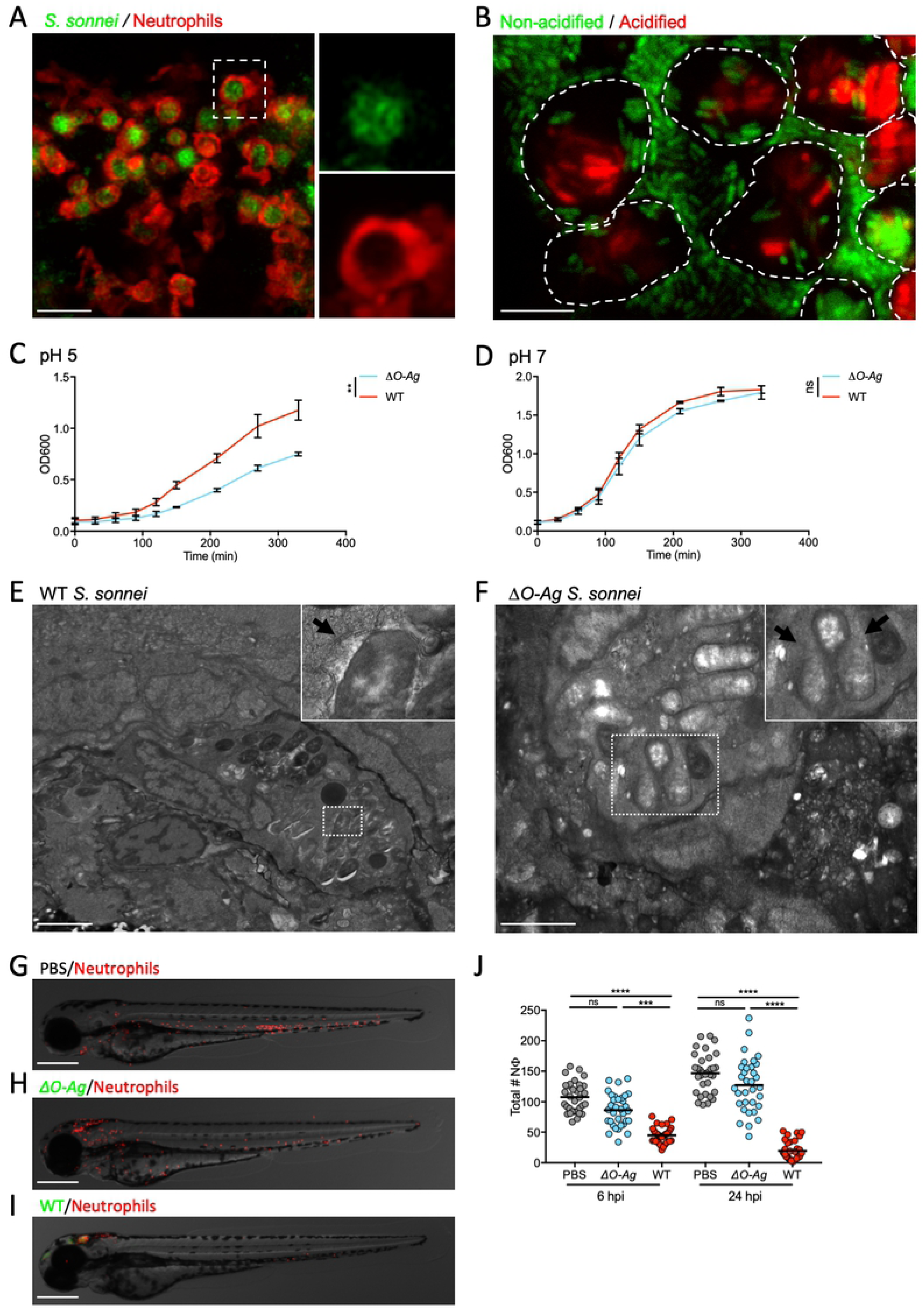
*S. sonnei* can resist phagolysosome acidification and promote neutrophil cell death in an O-antigen-dependent manner. **A. *S. sonnei* is collected by neutrophils in large phagosomes.** Larvae of the *Tg(lyz:dsRed)^nz50^* strain (labelling neutrophils) were injected in the HBV with WT GFP-*S. sonnei.* Image was taken at 3 hpi. Scale bar = 50 μm. **B. *S. sonnei* is acidified in immune cell phagosomes.** Larvae were injected in the HBV with WT GFP-*S. sonnei* which was also stained with pHrodo, a pH-sensitive dye that turns red in acidic environments. Dashed lines highlight the outline of individual phagocytes. Image taken at 4 hpi. Scale bar = 20 μm. **C,D. *S. sonnei* O-antigen contributes to acid tolerance *in vitro***. Growth curves of *ΔO-Ag* (blue) or WT (red) *S. sonnei,* cultured in tryptic soy broth adjusted to pH = 5 (C) or 7 (D). Statistics: unpaired t-test at the latest timepoint; **p<0.0021. **E,F. *S. sonnei* requires the O-antigen to survive in phagosomes.** Transmission electron micrographs of infected phagocytes from zebrafish larvae at 3 hpi with WT (E) or with *ΔO-Ag* (F) *S. sonnei*. E shows an intact phagocyte and *S. sonnei* residing within a phagosome (arrow points at phagosomal membrane). F shows that *ΔO-Ag S. sonnei* bacteria being degraded by a phagocyte (arrows point at region of major loss of bacterial cell integrity). Scale bars = 3 μm (E); 2 μm (F). **G-J. The O-antigen is required for *S. sonnei*-mediated killing of neutrophils.** Representative micrographs of larvae of the *Tg(lyz:dsRed)^nz50^* strain injected in the HBV with PBS (G), GFP-*ΔO-Ag* (H) or WT (I) *S. sonnei* at 6 hpi and quantification of total neutrophil number at 6 and 24 hpi (J). Statistics: Kruskal-Wallis test with Dunn’s correction; ns, non-significant; **p<0.0021; ****p<0.0001. Scale bars = 250 μm.

To investigate whether neutrophil cell death mediated by *S. sonnei* is dependent on O-Ag, we quantified neutrophils at the whole animal level in WT or *ΔO-Ag S. sonnei* infected larvae at 6 and 24 hpi. Infection with WT *S. sonnei* resulted in significantly more neutrophil cell death than infection with *ΔO-Ag S. sonnei* (**Fig. 5G-J)**. In the case of *S. flexneri* infection, neutrophils are recognised to die via necrosis [28]. Since no pharmacological reagent exists to directly test necrosis *in vivo*, we sought to rule out other cell death pathways in *S. sonnei* infected larvae and inhibited apoptosis, pyroptosis and/or necroptosis using the pan-caspase inhibitor Q-VD-OPh (an inhibitor of apoptosis and pyroptosis), Necrostatin-1 and/or Necrostatin-5 (inhibitors of necroptosis) (**Fig. S5E**). All inhibitors tested fail to significantly increase zebrafish survival. Considering this, and that infected zebrafish cells appeared necrotic by TEM (**Fig. S5D**), we conclude that neutrophils infected with *S. sonnei* undergo necrosis (and not a programmed mechanism of cell death) because of bacterial survival enabled by *S. sonnei* O-Ag.

### Phagolysosome acidification controls *S. sonnei* clearance by zebrafish and human neutrophils

To test the role of phagolysosome acidification during *S. sonnei* infection *in vivo*, we treated infected larvae with bafilomycin, an inhibitor of vacuolar H^+^ ATPase (V-ATPase). Consistent with a role for phagolysosome acidification in *S. sonnei* control, bafilomycin treatment significantly increased zebrafish susceptibility to *S. sonnei* (**Fig. 6A**). Bafilomycin treatment also increased zebrafish susceptibility to *ΔO-Ag S. sonnei* (**Fig. 6B**), highlighting the virulence of *ΔO-Ag S. sonnei* in the absence of phagolysosome acidification.

**Figure 6.**
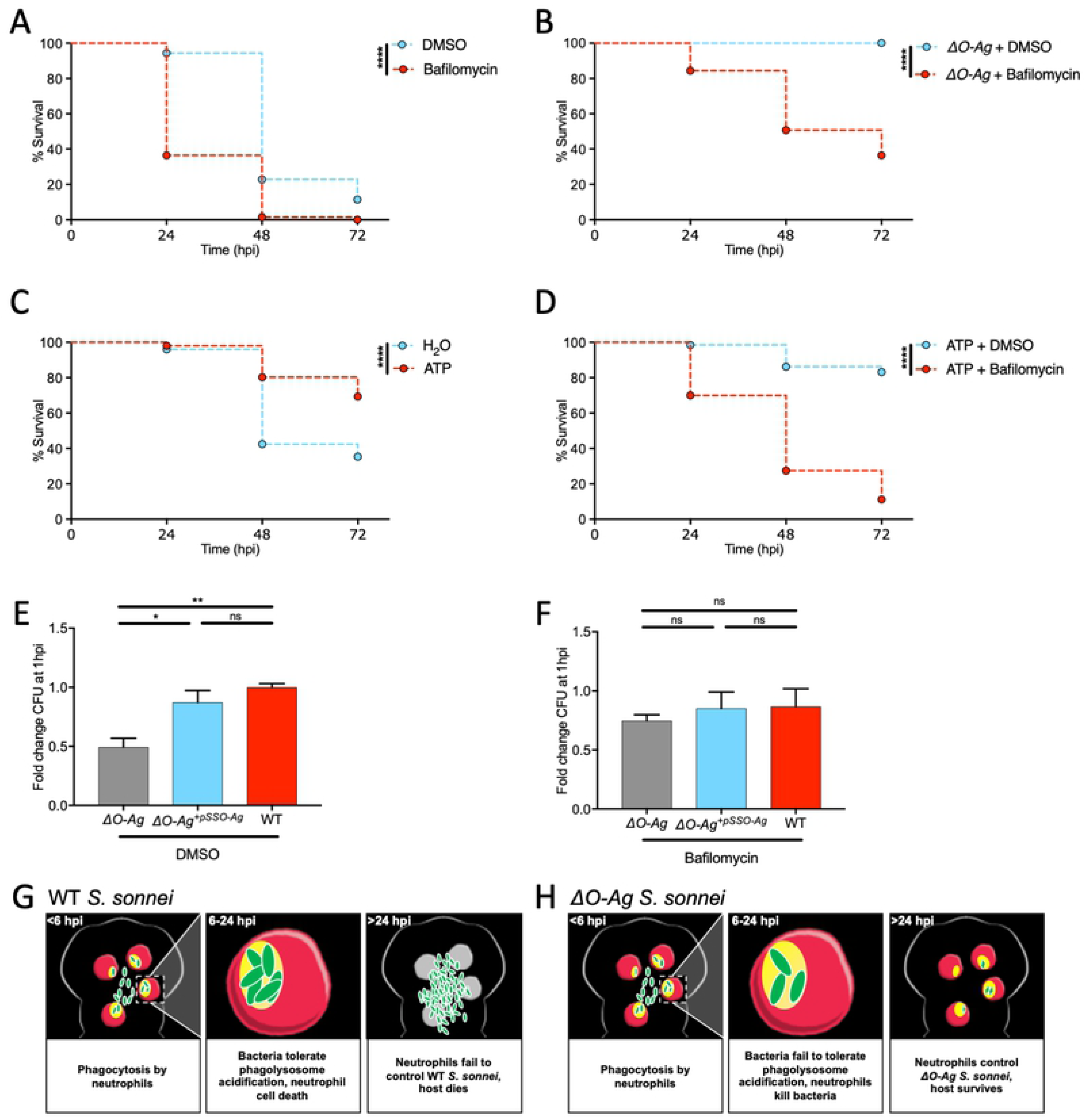
Phagolysosome acidification controls *S. sonnei* clearance by zebrafish and human neutrophils. **A,B. Bafilomycin treatment increases susceptibility to WT and *ΔO-Ag S. sonnei***. Survival curves of larvae treated with control DMSO vehicle (blue) or bafilomycin (red) upon infection in the HBV with WT (A) or *ΔO-Ag* (B) *S. sonnei*. Experiments are cumulative of 3 biological replicates. Bacterial input: ∼7000 CFU. Statistics: Log-rank (Mantel-Cox) test; ****p<0.0001. **C. ATP injections protect against *S. sonnei* infection.** Survival curves of larvae injected in the HBV with control water (blue) or ATP (red) 3 hours prior to infection of the same compartment with *S. sonnei*. Experiments are cumulative of 3 biological replicates. Bacterial input: ∼7000 CFU. Statistics: Log-rank (Mantel-Cox) test; ****p<0.0001. **D. Bafilomycin treatment and ATP injections counteract each other.** Survival curves of larvae injected in the HBV with ATP 3 hours prior to infection of the same compartment with *S. sonnei* and treatment with control DMSO vehicle (blue) or bafilomycin (red). Experiments are cumulative of 3 biological replicates. Bacterial input: ∼7000 CFU. Statistics: Log-rank (Mantel-Cox) test; ****p<0.0001. **E,F. *S. sonnei* O-Ag is required to counteract acidification-mediated clearance by human neutrophils.** *ΔO-Ag* (grey), complemenented strain (*ΔO-Ag^+pSSO-Ag^*, blue) or WT (red) *S. sonnei* were incubated with peripheral human neutrophils and exposed to DMSO (vehicle control treatment, E) or Bafilomycin (F). Difference in bacterial killing was quantified by plating from lysates of infected neutrophils at 1 hpi. Experiments are cumulative of 3 biological replicates from 3 independent donors. Statistics: one-way ANOVA with Sidak’s correction; ns, non-significant; *p<0.0332; **p<0.0021. **G,H. Model of *S. sonnei* O-antigen counteracting neutrophils *in vivo*.** Upon phagocytosis of WT *S. sonnei* (green), zebrafish neutrophils (red) rapidly acidify phagolysosomes containing bacteria. However, *S. sonnei* can tolerate this environment because of its O-antigen. *S. sonnei* replication leads to neutrophil and host death (G). *S. sonnei* without O-antigen fail to counteract acidification of neutrophil phagolysosomes. In this case, neutrophils clear infection and the host survives (H).

Considering that inhibition of phagolysosome acidification increases zebrafish susceptibility to *S. sonnei*, we hypothesised that promotion of acidification may overcome tolerance provided by O-Ag *in vivo*. V-ATPases mediate phagolysosome acidification by using ATP to pump protons in acidifying compartments. Injections of 200 μM ATP 3 h prior to *S. sonnei* infection significantly increases zebrafish survival (**Fig. 6C**). Bafilomycin directly antagonises V-ATPase activity. In agreement with this, treatment of *S. sonnei* infected larvae with bafilomycin counteracts the beneficial effects of ATP injection for host defence (**Fig. 6D**).

To test the role of *S. sonnei* O-Ag in human infection, we isolated peripheral neutrophils from healthy donors and infected them with WT or *ΔO-Ag S. sonnei* (**Fig. 6E**). In agreement with results from zebrafish infection, *ΔO-Ag S. sonnei* are significantly more susceptible to human neutrophil-mediated clearance than WT bacteria and bafilomycin treatment increased susceptibility of human neutrophils to *ΔO-Ag S. sonnei* (**Fig. 6F**). Plasmid reintroduction of the *O-Ag* biosynthesis system in *ΔO-Ag S. sonnei* (*ΔO-Ag+^pSSO-Ag^*) could restore the resistance of mutant bacteria to neutrophil killing to levels observed using WT bacteria (**Fig. 6E-F**). Collectively, these results show that *S. sonnei* O-Ag enables neutrophil tolerance in zebrafish and human neutrophils, and suggest that promotion of phagolysosome acidification is a novel approach to counteract *S. sonnei* infection.

### The innate immune system can be trained to control *S. sonnei in vivo*

Neutrophils of zebrafish larvae can be trained to protect against *S. flexneri* infection [24]. To test if we can enhance innate immunity to *S. sonnei*, we developed a *S. sonnei* reinfection assay (**Fig. 7A**). For this, larvae at 2 dpf were injected in the HBV with PBS or a sublethal dose (∼80 CFU) of WT or *ΔO-Ag* GFP-*S. sonnei*. At 48 hpi, we screened larvae and found that ∼20% of WT *S. sonnei* infected larvae are unable to clear infection (**Fig. S6A**); these larvae were therefore excluded from further analysis. Next, PBS-injected larvae or larvae clearing the primary infection (as determined by fluorescence microscopy) were infected with a lethal dose (∼8000 CFU) of mCherry-*S. sonnei*. Strikingly, injection of larvae with WT *S. sonnei* (but not *ΔO-Ag S. sonnei*) significantly increased survival, as compared to PBS-injected larvae (**Fig. 7B-C**). These experiments show that larvae exposed to a sublethal dose of *S. sonnei* are protected against a secondary lethal dose of *S. sonnei* in an O-Ag-dependent manner, and may have important implications in vaccine design.

**Figure 7.**
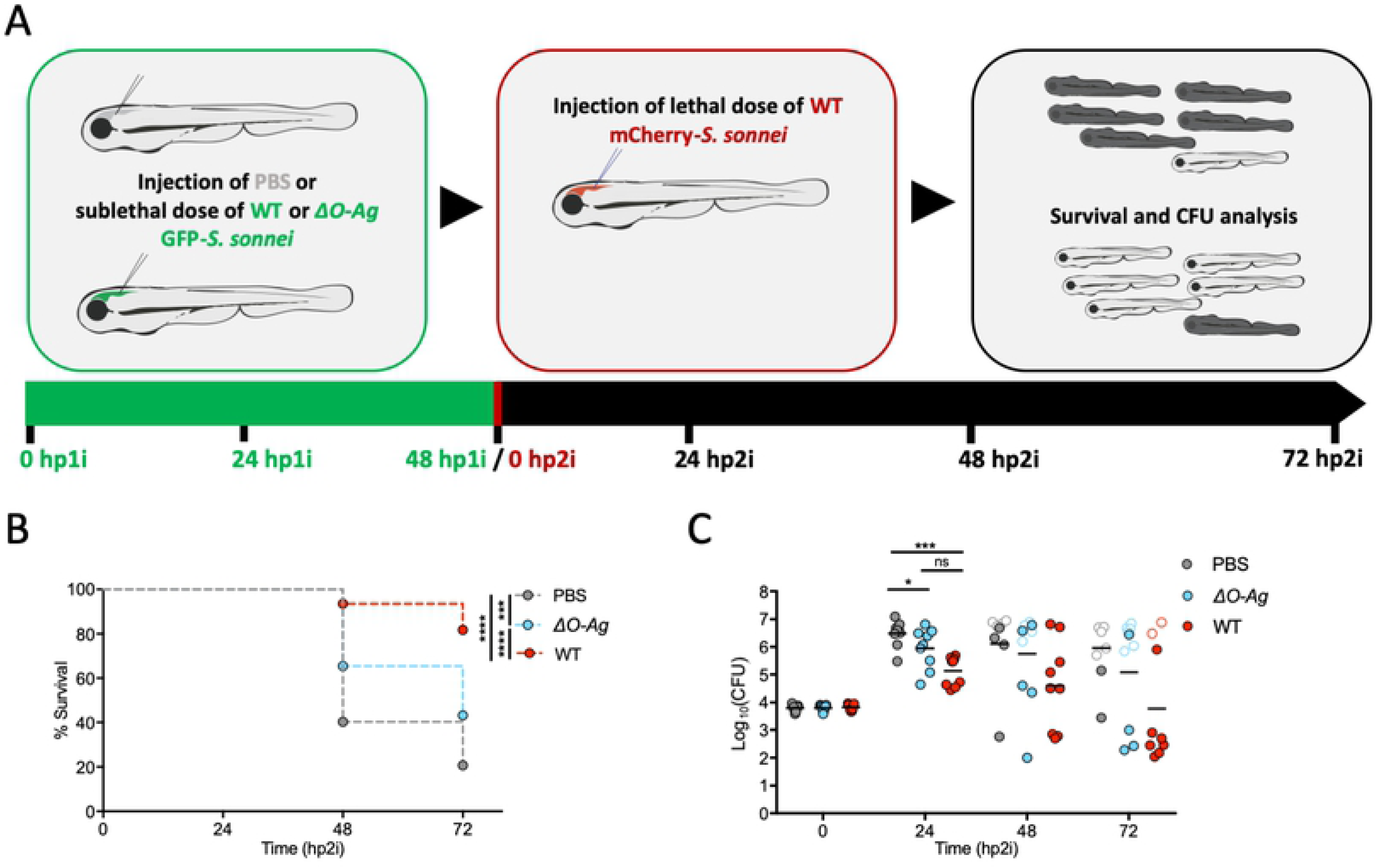
Innate immunity can be trained to control *S. sonnei in vivo*. **A. Model for *S. sonnei* reinfection**. 2 dpf embryos were primed with PBS or a sublethal dose of *S. sonnei*. At 48 hours post primary infection (hp1i), larvae where challenged with a lethal dose of *S. sonnei* and monitored by survival assay for 72 hours post-secondary infection (hp2i). **B,C. Innate immune training to *S. sonnei* is dependent on O-antigen.** Survival curves (B) and Log_10_-transformed CFU counts (C) of 4 dpf larvae infected in the HBV with lethal dose (∼8000 CFU) of mCherry-WT *S. sonnei.* 48 hours prior to infection with lethal dose, embryos were primed by delivering PBS (grey), or a sublethal dose (∼80 CFU) of GFP-*ΔO-Ag* (blue) or GFP-WT (red) *S. sonnei*. Experiments are cumulative of 4 (B) or 3 (C) biological replicates. In C, full symbols represent live larvae and empty symbols represent larvae that at the plating timepoint had died within the last 16 hours. Statistics: Log-rank (Mantel-Cox) test (G);); one-way ANOVA with Sidak’s correction on Log_10_-transformed data (H); ns, non-significant; *p<0.0332; ***p<0.0002; ****p<0.0001.

## Discussion

Why *S. sonnei* is emerging globally as a primary agent of bacillary dysentery has been unknown. Here, we discover that *S. sonnei* is more virulent than *S. flexneri in vivo* because of neutrophil tolerance mediated by its O-Ag. We also show that increased phagolysosomal acidification or innate immune training can promote *S. sonnei* clearance by neutrophils *in vivo* and propose new approaches to *S. sonnei* control.

The O-Ag, a lipopolysaccharide component of Gram-negative bacteria consisting of repetitive surface oligosaccharide units, is a major target for the immune system and bacteriophages. As a result, it is viewed that co-evolution of bacteria with their hosts or phages has led to significant variation in O-Ag structure/composition across bacterial strains [29]. In the case of most *Shigella spp*, significant variation of their *E. coli*-like O-Ag is observed across strains [27]. Interestingly, genes involved in *S. sonnei* O-Ag biosynthesis are non-homologous to those of other *Shigella spp*, highly conserved across *S. sonnei* strains and the only example of a virulence-plasmid encoded O-Ag system (in other *Shigella spp* the O-Ag is encoded by chromosomal genes) [27]. Consistent with this, acquisition of O-Ag genes from *P. shigelloides* is considered a defining event for the emergence of *S. sonnei* [15, 30]. We show that *S. sonnei* O-Ag enables bacteria to resist phagolysosome acidification and promotes neutrophil cell death. In agreement with this, chemical inhibition or promotion of phagolysosome acidification respectively decreases and increases neutrophil control of *S. sonnei* and zebrafish survival. In future studies it will be important to investigate the precise role of *S. sonnei* O-Ag in tolerance to phagolysosome acidification in neutrophils, and inspire new approaches for *S. sonnei* control.

Innate immune memory is a primitive form of immune memory conserved across vertebrates [31, 32]. We reveal that larvae injected with a sublethal dose of *S. sonnei* are protected against a secondary lethal dose of *S. sonnei* in an O-Ag-dependent manner. Although there is no vaccine currently available for *S. sonnei*, our results suggest that training innate immune memory against O-Ag should be considered for vaccine development. Moreover, it is tempting to speculate that innate immune memory may help to explain the increasing *S. sonnei* burden in regions where improved water sanitisation has eliminated *P. shigelloides* and subsequently reduced cross-immunisation against *S. sonnei* O-Ag [15, 16]. Consistent with this, the incidence of *S. sonnei* infection is mostly observed in very young children (<5 years old) [33–37], an age group where trained innate immunity has been shown to play an important protective role [31,32, 38–40].

Collectively, these findings reveal O-antigen as an important therapeutic target against bacillary dysentery. These findings also have major implications for our evolutionary understanding of *Shigella* and may explain the increasing burden of *S. sonnei* in developing countries.

## Materials and Methods

### Ethics statements

Animal experiments were performed according to the Animals (Scientific Procedures) Act 1986 and approved by the Home Office (Project licenses: PPL P84A89400 and P4E664E3C). All experiments were conducted up to 7 days post fertilisation.

Tissue samples from anonymised human donors (neutrophils) were provided by the Imperial College Healthcare NHS Trust Tissue Bank 12275. Other investigators may have received samples from these same tissues.

### Zebrafish

Zebrafish lines used here were the wildtype (WT) AB strain, macrophage reporter line *Tg(mpeg1:Gal4-FF)^gl25^/Tg(UAS-E1b:nfsB.mCherry)^c264^* and neutrophil reporter line *Tg(lyz:dsRed)^nz50^*. Unless specified otherwise, eggs, embryos and larvae were reared at 28.5°C in Petri dishes containing embryo medium, consisting of 0.5x E2 water supplemented with 0.3 μg/ml methylene blue (Sigma-Aldrich, St. Louis, Missouri). For injections and *in vivo* imaging, anaesthesia was obtained with buffered 200 μg/ml tricaine (Sigma-Aldrich) in embryo medium. Protocols are in compliance with standard procedures as reported at zfin.org.

### Bacterial preparation and infection delivery

Unless specified otherwise, GFP fluorescent or non-fluorescent *S. flexneri* M90T or *S. sonnei* 53G were used. Mutant, transgenic and WT strains are as indicated in the Figure legends and further detailed in **Table S3**.

Bacteria were grown on trypticase soy agar (TSA, Sigma-Aldrich) plates containing 0.01% Congo red (Sigma-Aldrich) supplemented, when appropriate, with antibiotics (Carbenicillin 100 μg/ml (Sigma-Aldrich), Kanamycin 50 μg/ml (Sigma-Aldrich), Streptomycin 50 μg/ml (Sigma-Aldrich)). Individual colonies were selected and grown O/N, 37°C/200 rpm, in 5 ml trypticase soy broth (TSB, Sigma-Aldrich) supplemented with the appropriate antibiotics as above. For injections, bacteria were grown to Log phase by diluting 400 μl of O/N culture in 20 ml of fresh TSB (supplemented, where appropriate, with 25 μg/ml Carbenicillin) and culturing as above until an optical density (OD) of 0.55-0.65 at 600 nm.

Bacteria were spun down, washed in phosphate buffer saline (PBS, Sigma-Aldrich) and resuspended at the desired concentration in a final injection buffer containing 2% polyvinylpyrrolidone (Sigma-Aldrich) and 0.5% phenol red (Sigma-Aldrich) in PBS (injection buffer alone is referred into the text as PBS group).

Unless specified otherwise, 1-2 nl of bacterial suspension (bacterial load as indicated in the individual experiments) or control solution were microinjected in the hindbrain ventricle (HBV) of 3 days post-fertilisation (dpf) zebrafish larvae (or at 2 dpf followed by reinfection at 4 dpf for reinfection assays, **Fig. 7B,C**). In **Fig. S1G,H** infection was delivered intravenously (IV), via the Duct of Cuvier.

Bacterial enumeration was performed *a posteriori* by mechanical disruption of infected larvae in 0.4% Triton X-100 (Sigma-Aldrich) and plating of serial dilutions onto Congo red-TSA plates. The *S. sonnei* 53G *Δtssb* strain was created using the λ Red recombinase approach [41]. Briefly, a Kan^R^ construct with flanking extensions homologous to the upstream and downstream regions of *tssB* was created by overlapping PCR using primers #1 and #2 (upstream *tssB*), #3 and #4 (downstream *tssB*) and #5 and #6 (Kan^R^). Primer sequences are reported in **Table S4**. The linear fragment was amplified (primers #1 and #4), parental plasmid removed by DpnI digestion and the linear fragment gel purified. The linear fragment was transformed into electrocompetent *S. sonnei* 53G containing pKD46; the λ Red recombinase genes on this plasmid were induced by arabinose prior to transformation. *Δtssb* colonies were selected on Kanamycin plates, and gene disruption was verified by multiple PCRs (primers #7 and #8, #7 and #9). pKD46 loss was confirmed by Ampicillin sensitivity. Successful gene disruption was also confirmed by sequencing of the entire region using primers #9 and #10.

To quantify the loss of virulence plasmid (**Fig. S3E**) GFP-labeled bacteria were injected in zebrafish larvae (1nl, ∼7000 CFU/nl) as above or spotted onto Congo red-TSA plates (10 μl, ∼7000 CFU/nl). Larvae and plates were incubated at 28.5°C for 24 h, bacteria were harvested from plates or larvae, and plated in serial dilutions on Carbenicillin-supplemented plates, grown O/N at 37°C and quantified for loss of virulence plasmid (white versus red colonies) as described elsewhere [26].

To address sensitivity of bacterial strains to acidic pH (**Fig. 5C,D, Fig. S5A,B**), bacteria were first grown O/N as described above, diluted to the same initial OD_600_ (0.10) in pH-adjusted TSB and bacterial growth was monitored by OD_600_ measurements at different timepoints. The pH of TSB was adjusted by addition of few drops of concentrated HCl (Sigma-Aldrich).

### Infection of human neutrophils

At least 5 ml of peripheral blood was drawn from healthy volunteers, using EDTA as an anticoagulant (BD Vacutainer, Becton Dickinson, Franklin Lakes, New Jersey). Neutrophils were isolated by gradient centrifugation using Polymorphprep^TM^ (Axis-Shield, Dundee, UK), according to the manufacturer’s guidelines and previously described protocols [42]. Residual erythrocytes were removed by incubation for 10 minutes at 37°C in erythrocyte lysis buffer, consisting of 0.83% w/v NH_4_Cl (Sigma-Aldrich), 10 mM NaOH-buffered HEPES (4-(2-hydroxyethyl)-1-piperazineethanesulfonic acid, Sigma-Aldrich), pH 7.4. Purified neutrophils were washed in Hank’s Balanced Salt Solution (HBSS) without Calcium and Magnesium (HBSS -Ca^2+^/-Mg^2+^, Thermofisher scientific, Waltham, Massachusetts), resuspended in neutrophil medium, consisting of HBSS with Calcium and Magnesium (HBSS +Ca^2+^/+Mg^2+^, Thermofisher scientific) and 0.1% porcine gelatin (Sigma-Aldrich), counted using trypan blue staining, and ultimately diluted at a density of 2 x 10^6^ live cells/ml in neutrophil medium [42]. Prior receiving infection, 10^5^ neutrophils (50 μl of neutrophil resuspension) were pre-incubated with Bafilomycin (111,11 nM, Sigma-Aldrich) [43, 44] or DMSO at vehicle control levels for 30 minutes with gentle shaking at 37°C in 48-well plates and in a total volume of 180 μl of neutrophil medium.

For neutrophil infections, *Shigella* was cultured as described above, but ultimately resuspended in neutrophil medium at a density of 5 x 10^4^ CFU/ml and 10^3^ bacteria (20 μl of bacterial resuspension) were added to the neutrophil resuspension and incubated for 1 h with gentle shaking at 37°C. Neutrophils were lysed by incubation on ice and addition of 7.5 μl of 0.4% Triton X-100 (Sigma-Aldrich) per well. Total CFUs were calculated by plating 20 μl of the lysate, comparing infected neutrophil samples to control samples, lacking neutrophils [42].

### pHrodo staining

pHrodo™ Red, succinimidyl ester (Thermofisher scientific) was prepared according to the manufacturer’s guidelines. 0.25 μl of stock solution were used to stain 200 μl of a ∼7000 CFU/nl bacterial suspension in PBS. Bacteria were incubated in the dark at 28.5°C for 30 minutes, washed 3 times in PBS, resuspended in 2% polyvinylpyrrolidone and 0.5% phenol red in PBS and injected in the HBV as above.

### Light and electron microscopy imaging

Stereo fluorescent microscopy images where acquired using Leica M205FA stereo fluorescent microscopes (Leica, Wetzlar, Germany). Zebrafsh larvae were anesthetised and aligned on 1% agarose plates in embryo medium.

For high-resolution confocal microscopy, imaging was performed using a Zeiss LSM 880 (Carl Zeiss, Oberkochen, Germany). Larvae were positioned in 35-mm-diameter glass-bottom MatTek dishes and imaged with 20× air or 40× water immersion objectives. Image files were processed using ImageJ/FIJI software.

For electron microscopy analysis, infected zebrafish larvae and controls were fixed at 3 hpi in 0.5% glutaraldehyde/200 nM sodium cacodylate buffer for 2 h and washed in cacodylate buffer only. Samples where then fixed in reduced 1% osmium tetroxide/1.5% potassium ferricyanide for 60 min, washed in distilled water and stained overnight at 4°C in 0.5% magnesium uranyl acetate. Specimens were then washed in distilled water, dehydrated in graded ethanol, infiltrated with propylene oxide and then graded Epon/PO mixtures until final embedding in full Epon resin in coffin moulds. Resin was allowed to polymerise at 56°C overnight, then semi-thin survey sections were cut and stained. Final ultrathin sections (typically 50–70 nm) and serial sections were collected on Formvar coated slot grids, stained with Reynold’s lead citrate and examined in a FEI Tecnai electron microscope with CCD camera image acquisition system.

### qRT-PCR

qRT-PCRs were performed using StepOne Plus machine (Applied Biosystems, Foster City, California) and a SYBR green master mix (Applied Biosystems). Briefly, RNA was isolated from pools of whole larvae using RNeasy mini kit (Qiagen, Hilden, Germany). cDNA was obtained using a QuantiTect reverse transcription kit (Qiagen). Samples were run in technical duplicates and quantification was obtained using the 2*^−ΔΔ^*^CT^ method and *eef1a1a* as a housekeeping gene. **Table S4** reports all primers used in this study.

### Enumeration of immune cells

For recruitment assays, immune cells attracted to the infection site were enumerated from images by counting *Tg(mpeg1:Gal4-FF)^gl25^/Tg(UAS-E1b:nfsB.mCherry)^c264^* (for macrophages) or *Tg(lyz:dsRed)^nz50^* (for neutrophils) positive cells in the hindbrain/midbrain. To quantify neutrophil death, the same neutrophil line was used to count immune cells at the whole animal level.

### Chemical treatments, ablations and knockdowns in zebrafish

Macrophages were ablated by exposing hatched 2 dpf *Tg(mpeg1:Gal4-FF)^gl25^/Tg(UAS-E1b:nfsB.mCherry)^c264^* embryos to Metronidazole (Mtz, Sigma-Aldrich) at 100 μM concentration in 1% DMSO (Sigma-Aldrich) for 24 h [45]. Treatment with 1% DMSO alone was used as a control.

Morpholinos were purchased from Gene Tools (Philomath, Oregon). Injections of *pu.1* (*spi1ab*) morpholino (2 nl, 0.5 mM in 0.5% phenol red) were performed at 1-cell stage and the same volume and concentration of a standard control morpholino was used as negative control. Morpholino sequences are reported in **Table S4**.

Necrostatin-1 (10 μM, Santa Cruz Biotechnology, Dallas, Texas), Necrostatin-5 (10 μM, Santa Cruz) [46], Q-VD-OPh (50 μM, Sigma-Aldrich) [47] and Bafilomycin (200 nM, Sigma-Aldrich) [43, 48] were provided by bath exposure from 0 hpi for the whole infection course. Exposure to DMSO at vehicle concentration (0.67% for Necrostatin-1, Necrostatin-5, Q-VD-OPh, and 0.2% for Bafilomycin) was used as a control. Priming with ATP injections was performed by injecting 1 nl of 200 mM ATP in the hindbrain 3 h prior infection. Injection of 1 nl of sterile water was used as a vehicle control.

### Dual-RNAseq sample preparation and analysis

RNA samples for dual-RNAseq were extracted in triplicate from infected larvae at 24 hpi using 24 larvae/sample. As a control for the host transcriptome, RNA was isolated from corresponding PBS injected larvae at the same timepoint. As a control for *S. sonnei* transcriptome, RNA was isolated from the same culture used for injection, but diluted 50x and subcultured at 28.5°C until it reached the OD of ∼0.6 in a total volume of 5 ml. Samples were snap frozen at -80°C, then 100 μl of RNA protect bacteria reagent (Qiagen) was added, followed by mechanical trituration with a pestle blender. Samples were supplemented with 100 μl of 30 mM Tris-HCl/1 mM EDTA solution at pH = 8, 33 μl of 50 mg/ml lysozyme (Thermo Scientific), 33 μl of proteinase K >600 U/ml (Thermo Scientific) and shaken for 20 min RT. Lysis was completed by adding 700 μl of RTL buffer (Qiagen), 3 μl of 1 M dithiothreitol (Sigma-Aldrich) and mechanical disruption. Undigested debris were spun down 3 min 10000 rpm and the supernatant was supplemented with 500 μl of 100% ethanol prior loading onto RNeasy mini columns (Qiagen). From this step onwards, the manufacturer’s guidelines were followed for RNA purification. RNA quality and integrity were assessed by using NanoDrop and Non-denaturing agarose gel electrophoresis. For further quality check, RNA sequencing, library construction and reads count, samples were outsourced to Vertis Biotechnologie AG (Freising, Germany). Bacterial and host mRNA were enriched prior library preparation by using Ribo-Zero Gold rRNA Removal Kit (Epidemiology, Illumina, San Diego, California). The zebrafish genome assembly GRCz11 (http://www.ensembl.org/Danio_rerio) and the *S. sonnei* 53G genome assembly ASM28371v1 (https://www.ncbi.nlm.nih.gov/assembly/406998) were used to guide mapping of host and pathogen reads, respectively. The average library depths for the different sample groups were: 8658688 +/- 598491 reads (PBS injected larvae), 7771779 +/- 804881 reads (infected larvae), 11315345 +/- 551602 reads (*S. sonnei in vitro*) and 702385 +/- 12785 reads (*S. sonnei in vivo*).

RNAseq statistical analysis was performed using DESeq2 package in R [49, 50]. Genes that were not represented with at least 6 reads cumulative from all samples were excluded from the analysis *a priori*. Genes were accepted as differentially expressed if the DESeq2 Log_2_(Fold Change) > |1| and DESeq2 adjusted p value (padj) < 0.05. Heatmaps (**Fig. 2D,E**) were obtained from counts per million (CPM) reads using the “pheatmap” package (https://CRAN.R-project.org/package=pheatmap). Principal component analysis (PCA) (**Fig. S2A,B**) was also performed in R from CPM reads, using the dedicated PCA tools [50]. All the other graphs were generated using GraphPad Prism 7 (GraphPad Software, San Diego, California). Host pathway enrichment analysis and enrichment of transcription factor binding sites were performed using ShinyGO v0.60 (http://bioinformatics.sdstate.edu/go/) [51]. Due to poor annotations of *S. sonnei* Gene Ontology functions, pathogen pathway enrichment analysis was performed manually on the top 50 differentially expressed genes, inferring protein funtions from *E. coli* homologues of *S. sonnei* genes. Homologues were identified by direct protein BLAST using the protein database available via uniprot.org.

### Statistical analysis and data processing

Except for graphs performed in R, all other graphs and statistical analyses were performed using GraphPad Prism 7. Statistical difference for survival curves were analysed using a Log-rank (Mantel-Cox) test. Differences in CFU recovery and gene expression levels were quantified on Log_10_-transformed or Log_2_-transformed data, respectively. To avoid Log(0), i.e., when no colonies were recovered, the CFU counts were assigned as 1. When only 2 groups were compared, significant differences were tested using an unpaired t-test at each timepoint. When more than 2 groups were compared, a one-way ANOVA with Sidak’s correction was used. Unpaired t-test (comparison between 2 groups) or one-way ANOVA with Sidak’s correction (comparison between more than 2 groups) was also applied to **Fig. S3E**, **Fig. 5C,D, Fig. S5A,B** and **Fig. 6. E,F** but on non-transformed data, as in this case a parametric distribution could be assumed. Statistics for categorical data were obtained by a two-sided *chi*-squared contingency test (**Fig. S1D, Fig. S6A**). For statistical quantification of immune cell numbers (non-parametric data), a two-tailed Mann-Whitney test (comparison between 2 groups) or a Kruskal-Wallis test with Dunn’s correction (comparison between more than 2 groups) was used.

## Acknowledgements

We thank Margarida C. Gomes and Nagisa Yoshida for helpful discussion.

## Supporting Information Legends

**Figure S1. (Related to Fig. 1) *S. sonnei* is more virulent than *S. flexneri* in a zebrafish infection model**

**A,B. Dose response to *S. sonnei* infection.** Survival curves (A) and Log_10_-transformed CFU counts (B) of larvae injected in the HBV with increasing doses of *S. sonnei.* ∼200 CFU range: 100-300 CFU (grey); ∼600 CFU range: 400-700 CFU (blue); ∼1500 CFU range: 1000-2000 CFU (green); ∼5000 CFU: 4000-6000 CFU (red). Experiments are cumulative of 3 biological replicates. In B, full symbols represent live larvae and empty symbols represent larvae that at the plating timepoint had died within the last 16 hours. Statistics: Log-rank (Mantel-Cox) test (A); one-way ANOVA with Sidak’s correction on Log_10_-transformed data (B); ns, non-significant; *p<0.0332; **p<0.0021; ****p<0.0001.

**C-F. *S. sonnei* can disseminate from the injection site**. Representative images of three larvae injected in the HBV with ∼20000 CFU of GFP-labelled *S. flexneri* (C, compare to **Fig. 1E,F**). The frequency of larvae with bacterial dissemination out of the HBV at 24 hpi is significantly higher for the *S. sonnei-*infected group when compared to the *S. flexneri* infected group (even when *S. flexneri* input is ∼3-fold higher than *S. sonnei* input) (D). Survival curves (E) and Log_10_-transformed CFU counts (F) of larvae injected in the HBV with ∼7000 CFU *S. flexneri* (grey), ∼20000 CFU of *S. flexneri* (blue) or ∼7000 CFU *S. sonnei* (red). Experiments are cumulative of 3 biological replicates. In E, full symbols represent live larvae and empty symbols represent larvae that at the plating timepoint had died within the last 16 hours. Statistics: two-sided *chi*-squared contingency test (D); Log-rank (Mantel-Cox) test (E); one-way ANOVA with Sidak’s correction on Log_10_-transformed data (F); ns, non-significant; **p<0.01; ****p<0.0001. Scale bar = 1 mm.

**G,H. *S. sonnei* is more virulent than *S. flexneri* in an intravenous infection model.** Survival curves (G) and Log_10_-transformed CFU counts (H) of larvae injected intravenously (IV, via the duct of Cuvier) with PBS (grey), *S. flexneri* (blue) or *S. sonnei* (red). Experiments are cumulative of 2 biological replicates. In H, full symbols represent live larvae and empty symbols represent larvae that at the plating timepoint had died within the last 16 hours. Statistics: Log-rank (Mantel-Cox) test (G); unpaired t-test on Log_10_-transformed data (H); ns, non-significant; ****p<0.0001.

**I,J. A clinical isolate of *S. sonnei* is more virulent than a clinical isolate of *S. flexneri.*** Survival curves (I) and Log_10_-transformed CFU counts (J) of larvae injected in the HBV with *S. flexneri* isolate 2457T (blue) or *S. sonnei* isolate 381 (red). Experiments are cumulative of 3 biological replicates. In J, full symbols represent live larvae and empty symbols represent larvae that at the plating timepoint had died within the last 16 hours. Statistics: Log-rank (Mantel-Cox) test (I); unpaired t-test on Log_10_-transformed data (J); **p<0.0021; ****p<0.0001.

**K-N. *S. sonnei* is more virulent than *S. flexneri* at 32.5°C and 37°C**. Survival curves (K,M) and Log_10_-transformed CFU counts (L,N) of larvae injected in the HBV with *S. flexneri* (blue) or *S. sonnei* (red) at 32.5°C (K,L) or at 37°C (M,N). Experiments are cumulative of 2 biological replicates. In L,N, full symbols represent live larvae and empty symbols represent larvae that at the plating timepoint had died within the last 16 hours. Statistics: Log-rank (Mantel-Cox) test (K,M); unpaired t-test on Log_10_-transformed data (L,N); ns, non-significant; *p<0.0332; ****p<0.0001.

**Figure S2. (Related to Fig. 2) Whole animal dual-RNAseq profiling of *S. sonnei* infected larvae**

**A,B. Principal component analysis (PCA) of *S. sonnei* and zebrafish larvae transcriptomes.** Analysis was performed on counts per million (CPM) reads values, using the dedicated PCA tools in R. Individual biological replicates (R1, R2, R3) for control (blue) and infected (red) conditions are reported. % in brackets indicate the variance of dimension explained by each principal component. Plot in A refers to *S. sonnei* genes and plot in B refers to zebrafish genes.

**C,D. Boxplots representing the distribution of reads within the RNAseq libraries of each individual sample.** Boxplots represent the sample median CPM reads with interquartile range, while whiskers indicate the 2.5-97.5 percentile range. Control samples are indicated in blue and infection samples are indicted in red. Biological replicates (R1, R2, R3) are also indicated. Plot in C refers to *S. sonnei* gene libraries and plot in D refers to zebrafish gene libraries.

**E-H. Distribution histograms of significantly differentially expressed genes in *S. sonnei* and zebrafish larvae during infections.** Each bar represents the number of significantly differentially expressed genes (repressed, blue (E,F); induced, red (G,H)) in each interval of Log_2_(FC). Plots in E,G refer to *S. sonnei* genes, while plots in F,H refer to zebrafish genes.

**I. Induction of well established inflammatory markers in the RNAseq transcriptome.** Bars indicate the average CPM reads for representative inflammatory marker. Compare to induction of same genes tested independently by qRT-PCR at the same timepoint in **Fig. 1D**. Statistics: unpaired t-test on Log_2_-transformed data; **p<0.0021; ***p<0.0002; ****p< 0.0001.

**J. Pathway enrichment analysis of *S. sonnei* during infection *in vivo*.** Pathway enrichment analysis was performed manually using the top 50 differentially expressed genes. Gene functions were inferred from *E. coli* annotations (accessible via uniprot.org). % are relative to all 50 genes analysed. A variety of stress response processes are induced in *S. sonnei in vivo* in the zebrafish larvae (i.e. amino acid metabolism, acid resistance systems, purine metabolism and DNA damage repair, regulation of transcription or translation and protection against oxidative stress). Upregulated genes involved in acidic adaptation include GadA, GadB, GadC and their transcriptional regulator GadE (the Gad pathway constitutes a glutamate-dependent system to maintain neutral cytoplasmic pH in acid conditions) [52], as well as HdeA, HdeB and HdeD (the Hde pathway constitutes an acid-induced chaperone system to preven protein misfolding in acid environment) [53]. In agreement with upregulation of acid resistance, 60% (9/15) of the genes associated with metabolic changes are specifically involved in the biosynthesis of glutamate (essential to fuel the GadA/B/C system) or arginine (shown to activate an arginine-dependent acid resistance system [54]).

**K. Pathway enrichment analysis of *S. sonnei*-infected zebrafish larve.** Pathway enrichment analysis was performed using ShinyGO v0.60 (http://bioinformatics.sdstate.edu/go/). % are relative to all the genes bioinformatically annotated to the pathway of interest. A variety of immune-related processes are induced in zebrafish larvae in response to *S. sonnei* infection, including leukocyte (especially neutrophil) chemotaxis, response to cytokines and inflammation.

**Figure S3. (Related to Fig. 3) *S. sonnei* virulence depends on its O-antigen**

**A,B. Schematic of *S. sonnei* and *S. flexneri***. Both *S. flexneri* and *S. sonnei* virulence plasmid encodes a type 3 secretion system (T3SS). However, differently than *S. flexneri, S. sonnei* virulence plasmid (pSS) encodes genes for the biosynthesis of a capsule and O-antigen (O-Ag) non-homologous to those of other *Escherichia* and *Shigella* species. *S. sonnei* additionally encodes a type 6 secretion system (T6SS) on the bacterial chromosome. The schematic in B also reports (in grey and between brackets) the name of the mutants used in the study.

**C,D. Virulence of *S. sonnei* in zebrafish does not depend on the T6SS or capsule.** Survival curves (C) and Log_10_-transformed CFU counts (D) of larvae injected in the HBV with *S. sonnei Δtssb* (grey), *Δg4c (blue),* or WT (red) strains. Experiments are cumulative of 4 (C) or 3 (D) biological replicates. In D, full symbols represent live larvae and empty symbols represent larvae that at the plating timepoint had died within the last 16 hours. Statistics: Log-rank (Mantel-Cox) test (C); one-way ANOVA with Sidak’s correction on Log_10_-transformed data (D); ns, non-significant.

**E. *S. sonnei* virulence plasmid is maintained *in vivo* in zebrafish at 28.5°C.** Larvae were injected with 1nl (∼7000 CFU) in the HBV with *S. flexneri* (blue) or *S. sonne*i (red) for 24 h at 28.5°C. Control bacteria (TSA plates) were spotted onto tryptic soy agar plates (10 μl of the bacterial inoculum/spot) and also grown for 24 h at 28.5°C. Bacteria were then harvested from larvae or plates and grown on Congo-Red plates at 37°C to quantify colonies that lost the virulence plasmid. Experiments are cumulative of 2 biological replicates. Statistics: one-way ANOVA with Sidak’s correction; ns, non-significant; *p<0.0332; **p<0.0021.

**F. Comparison of *S. sonnei* and *S. flexneri* O-antigen.** *S. sonnei* 53G O-antigen has a unique sugar composition compared to the O-antigen of *S. flexneri* M90T. Figure legend abbreviations: AN: 2-Acetamido-2-deoxy-L-altruronic acid (L-AltNAc); FN: 2-Acetamido-4-amino-2,4-dideoxy-D-fucose (D-FucNAc); GN: 2-Acetamido-2-deoxy-D-glucose (D-GlcNAc); R: L-Rhamnose (L-Rha); G: D-Glucose (D-Glc); Ac: O-acetyl. Adapted from [55].

**G. Induction of the O-antigen chain length determinant protein *wzzB in vivo*.** Dual-RNAseq profiling shows that the *wzzB* is significantly upregulated in *S. sonnei* infected zebrafish. Bars indicate the average CPM reads. Statistics: unpaired t-test on Log_2_-transformed data; ***p<0.0002.

**Figure S4. (related to Fig. 4) *S. sonnei* O-antigen can counteract clearance by zebrafish neutrophils**

**A-C. Chemical ablation of macrophages.** Representative micrographs (A,B) and quantification (C) of *Tg(mpeg1:Gal4-FF)^gl25^/Tg(UAS-E1b:nfsB.mCherry)^c264^* larvae which were treated with either Metronidazole (Mtz, macrophage ablated group, blue) or control DMSO vehicle (DMSO, red) prior to infection in the HBV with *S. sonnei*. Experiments are cumulative of 2 biological replicates. Scale bars = 250 μm.

**D,E. Macrophage ablation increases susceptibility to *S. flexneri*.** Survival curves (D) and Log_10_-transformed CFU counts (E) of *Tg(mpeg1:Gal4-FF)^gl25^/Tg(UAS-E1b:nfsB.mCherry)^c264^* larvae which were treated with either Metronidazole (Mtz, macrophage ablated group, blue) or control DMSO vehicle (DMSO, red) prior to infection in the HBV with *S. flexneri*. Experiments are cumulative of 3 biological replicates. In E, full symbols represent live larvae and empty symbols represent larvae that at the plating timepoint had died within the last 16 hours. Statistics: Log-rank (Mantel-Cox) test (D); unpaired t-test on Log_10_-transformed data (E); ns, non-significant; *p<0.0332.

**F,G. *pu.1* morpholino knockdown increases susceptibility to *S. sonnei* when infections are performed at 30 hpf.** Survival curves (F) and Log_10_-transformed CFU counts (G) of *pu.1* morphant (blue) or control (red) larvae infected in the HBV with WT *S. sonnei*. Experiments are cumulative of 3 biological replicates. In G, full symbols represent live larvae and empty symbols represent larvae that at the plating timepoint had died within the last 16 hours. To allow full ablation of immune cells by morpholino knockdown, infections are performed at 30 hpf. Statistics: Log-rank (Mantel-Cox) test (F); unpaired t-test on Log_10_-transformed data (G); ns, non-significant; ****p<0.0001.

**Figure S5. (related to Fig. 5) *S. sonnei* can resist phagolysosome acidification and promote neutrophil cell death in an O-antigen-dependent manner**

**A,B. *S. sonnei* O-antigen contributes to acid tolerance *in vitro***. Growth curves of *ΔO-Ag* (blue) or WT (red) *S. sonnei,* cultured in tryptic soy broth adjusted to pH = 4 (A) or 6 (B). Statistics: unpaired t-test at the latest timepoint; ns, non-significant.

**C. *S. sonnei* can replicate within phagosomes.** Transmission electron micrograph of an infected phagocyte from zebrafish larvae at 3 hpi with WT *S. sonnei*, showing a dividing *S. sonnei* cell. Scale bar = 4 μm.

**D.** *S. sonnei* infection of zebrafish cell promotes morphological features of necrosis. Transmission electron micrograph of an infected neutrophil from a zebrafish larva at 3 hpi with WT *S. sonnei*, showing signs of necrotic cell death (arrowheads point at area of extranuclear chromatin degradation). Scale bar = 4 μm.

**E. Pharmacological inhibition of necroptosis and/or apoptosis/pyroptosis does not protect zebrafish larvae.** Survival curves of larvae which were treated with Necrostatin-1 (grey), Necrostatin-5 (blue), Q-VD-OPh (green), Necrostatin-1 + Q-VD-OPh (red) or control DMSO vehicle (black) upon infection in the HBV with *S. sonnei*. Experiments are cumulative of 3 biological replicates. Bacterial input: ∼7000 CFU. Statistics: Log-rank (Mantel-Cox) test; ns, non-significant. Scale bar = 4 μm.

**Figure S6. (related to Fig. 7) Innate immunity can be trained to control *S. sonnei in vivo***

**A,B. Response of 2dpf zebrafish embryos to sublethal dose (∼80 CFU) of *S. sonnei*.** Approximately 80% of WT GFP-*S. sonnei* injected embryos (and 100% of *ΔO-Ag* GFP-*S. sonnei* control/clear infection (no detectable bacteria by fluorescence microscopy) by 48 hpi (A). Log_10_-transformed CFU counts from controller larvae (no detectable bacteria by fluorescence microscopy) infected in the HBV with GFP-*ΔO-Ag* (blue) or WT (red) *S. sonnei*. Prior to receiving the secondary lethal dose (∼8000 CFU) of mCherry-*S. sonnei*, ∼80% of WT GFP-*S. sonnei* injected controllers (and 100% of *ΔO-Ag* GFP-*S. sonnei* injected controllers) cleared the primary infection. Experiments are cumulative of 4 (A) or 3 (B) biological replicates. Statistics in A: two-sided *chi*-square contingency test.

**Table S1. Differentially expressed *S. sonnei* genes and pathogen pathway enrichment analysis *in vivo*.**

**Table S2. Differentially expressed zebrafish genes and host pathway enrichment analysis in response to *S. sonnei* infection.**

**Table S3. Bacterial strains used in this study.**

**Table S4. Primers and morpholinos used in this study.**

